# *PinMol*: Python application for designing molecular beacons for live cell imaging of endogenous mRNAs

**DOI:** 10.1101/294447

**Authors:** Livia V. Bayer, Omar S. Omar, Diana P. Bratu, Irina E. Catrina

## Abstract

Molecular beacons are nucleic acid oligomers labeled with a fluorophore and a quencher that fold in a hairpin-shaped structure, which fluoresce only when bound to their target RNA. They are used for the visualization of endogenous mRNAs in live cells. Here, we report a Python program (*PinMol*) that designs molecular beacons best suited for live cell imaging by using structural information from secondary structures of the target RNA, predicted via energy minimization approaches. *PinMol* takes into account the accessibility of the targeted regions, as well as the inter- and intramolecular interactions of each selected probe. To demonstrate its applicability, we synthesized an *oskar* mRNA-specific molecular beacon (osk1236), which is selected by *PinMol* to target a more accessible region than a manually designed *oskar*-specific molecular beacon (osk2216). We previously demonstrated osk2216 to be efficient in detecting *oskar* mRNA in *in vivo* experiments. Here, we show that osk1236 outperformed osk2216 in live cell imaging experiments.

## INTRODUCTION

Currently, there are a few methods that are employed to image RNA in live tissues and cells, and each has its advantages and disadvantages (Weil et al. 2010). We use the molecular beacon technology for live cell imaging of mRNA transport and localization in the fruit fly egg chamber (Bratu et al. 2003). Molecular beacons are oligonucleotides labeled with a fluorophore at the 5’ end and a quencher at the 3’ end that are designed to fold into a hairpin shape (Fig. 1). In this manuscript we use the terms “probe” and “stem” to refer to the loop or single stranded region of the hairpin with a sequence complementary to the mRNA target (18-26 nucleotides), and to the double-stranded region (five G/C rich base pairs), respectively. In the presence of target, the hybridization of the probe region with the target is energetically more favorable than preserving the duplex in the stem, which leads to the separation of the fluorophore/quencher pair and a significant increase in the fluorescence signal (Fig. 1). One major advantage of using this technology is that molecular beacons allow direct visualization of endogenous mRNAs. However, not many groups have reported *in vivo* mRNA imaging via molecular beacons. One reason is the difficulty in finding an efficient delivery method for the chosen live specimen. We have successfully introduced nuclease-resistant molecular beacons in fruit fly egg chambers via microinjection, to study transport and localization of endogenous maternal mRNAs (Bratu et al. 2003). Others have described the use of toxin-based membrane permeabilization and microporation for efficient cytoplasmic delivery of molecular beacons (Chen et al. 2011). More recently, gold-nanoparticles were shown to efficiently deliver molecular beacons into live breast cancer cells for multiplex detection of mRNAs (Jackson et al. 2016). Another reason for the limited *in vivo* use of molecular beacons is the challenge to design them, as several properties need to be optimized to ensure specific and efficient detection of the target RNA. Here, we describe a new open-source Python-based application that allows fast and easy prediction of molecular beacons tailored for live cell imaging.

**FIGURE 1.**
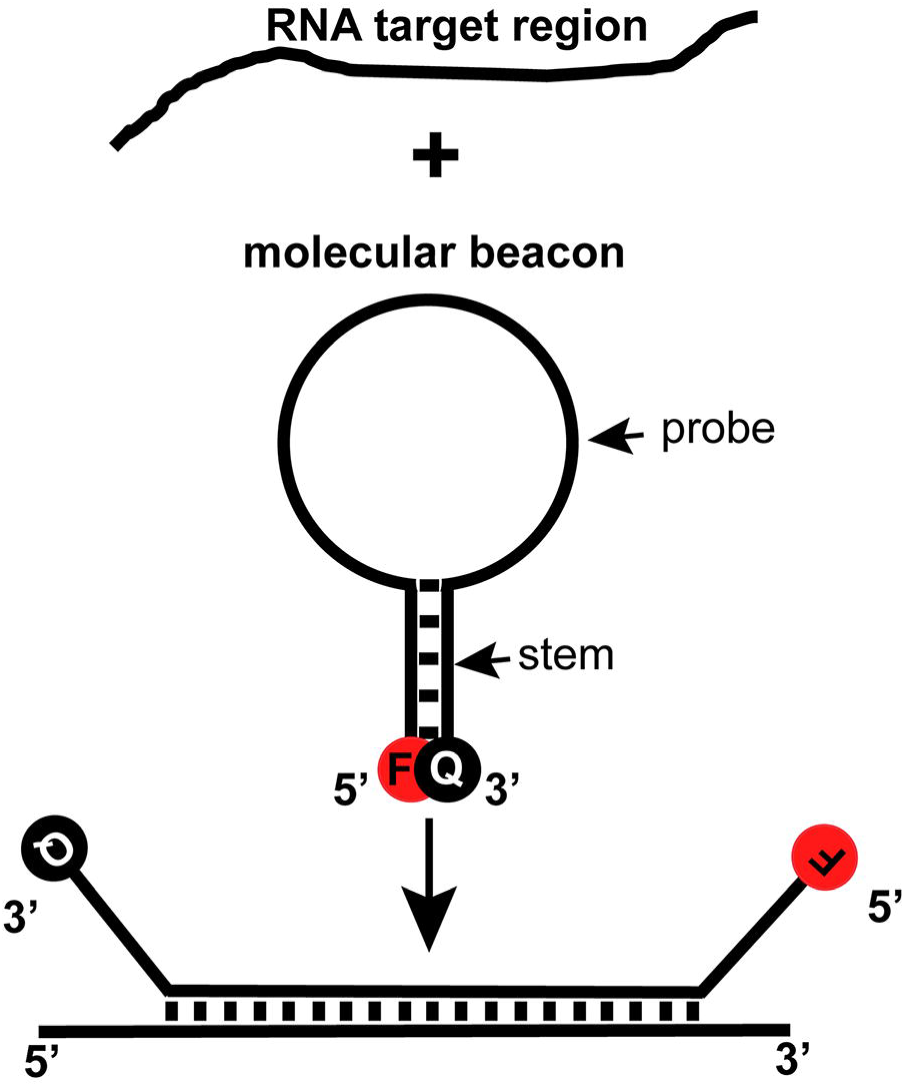
Molecular beacons, principle of operation. F = fluorophore, Q = quencher.

There are several stand-alone or server-based databases [*e.g.* miRBase (Griffiths-Jones et al. 2008)] and applications [*e.g.* TargetScan (Lewis et al. 2005), OligoWalk (Lu and Mathews 2008b), siRNA Wizard from InvivoGen, etc.] for the analysis and identification of targets of small RNAs, such as microRNAs (miRNA) and small interference RNAs (siRNA), and for the design of small probes that efficiently bind to mRNA targets. However, freely available programs for designing molecular beacons focus on their application for quantitative PCR (qPCR) primer design and analysis, and they usually limit the target region to less than 1,500 bases (*e.g.* OligoArchitect, from Sigma-Aldrich powered by Beacon Designer^™^, the allowed target length is 50-1,200 nt when template structure is taken into consideration) (Table 1).

**TABLE 1.**
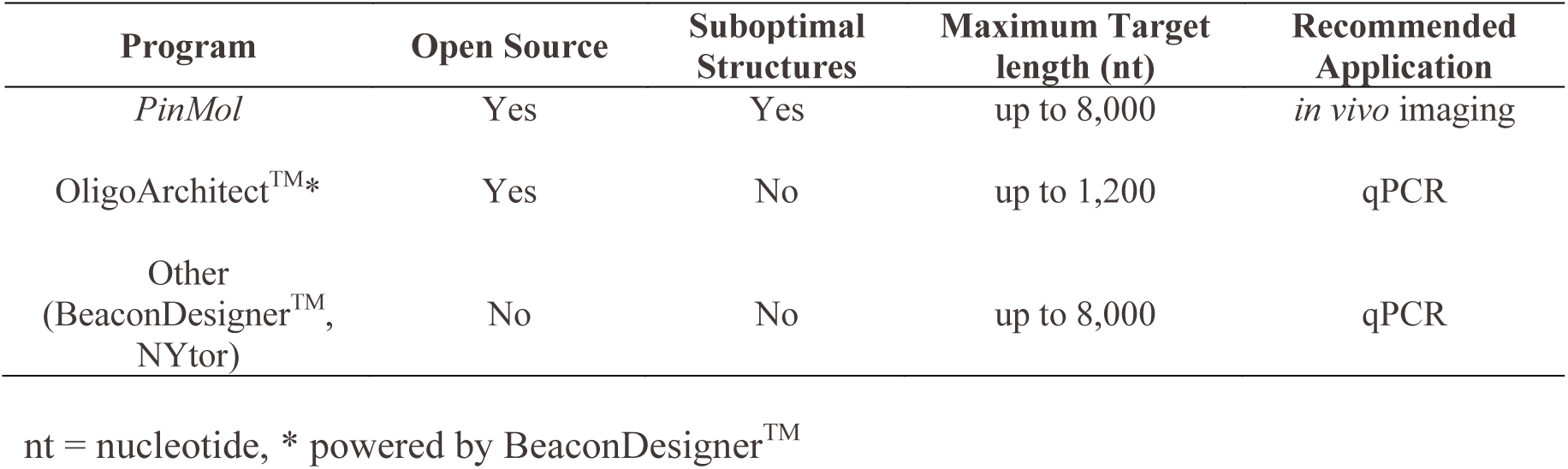
Comparison of programs for designing molecular beacons, which take into consideration the structure of the RNA target.

| Program | Open Source | Suboptimal Structures | Maximum Target length (nt) | Recommended Application |
| --- | --- | --- | --- | --- |
| *PinMol* | Yes | Yes | up to 8,000 | *in vivo* imaging |
| OligoArchitect<sup>TM</sup>* | Yes | No | up to 1,200 | qPCR |
| Other<br>(BeaconDesigner<sup>TM</sup>,<br>NYtor) | No | No | up to 8,000 | qPCR |

nt = nucleotide, \* powered by BeaconDesigner<sup>TM</sup>

*PinMol* (Python program for designing hair<u>Pin</u>-shaped <u>Mol</u>ecular beacons) permits the fast design of molecular beacons suited for live cell imaging by taking into consideration target accessibility to minimize false negative results, and cross homology to avoid false positive results.

## RESULTS

### The *PinMol* algorithm

#### Input File

To assess target-site accessibility, it is common to evaluate the predicted secondary structure of the RNA target of interest, which is generated using either phylogenetic analysis or free energy minimization algorithms. The efficiency of a candidate probe is directly correlated to the accuracy of the predicted secondary structure of the target RNA. When employing results from free energy minimization algorithms, it is recommended to consider the global structure of a target RNA rather than a local structure prediction of a limited region within that target (Lu and Mathews 2008a). *mfold*, a software commonly used to predict the secondary structure of nucleic acids using experimentally derived thermodynamic parameters, yields a minimum free energy (MFE) structure and a user-selected number of suboptimal (SO) structures (Zuker 2003). *mfold*, *RNAstructure* and the *Vienna* RNA*fold* package use similar algorithms, and although they use different sets of RNA thermodynamic parameters (*mfold*: 1999 Turner set; *RNAstructure*: 2004 Turner set; *Vienna* RNA*fold*: choice of 1999 or subset of 2004 Turner set, 2007 Andronescu set) that result in slightly different predictions, their performance of accuracy are comparable. *mfold* and *RNAstructure* predict MFE and suboptimal RNA secondary structures, while the RNA*fold* generates an MFE secondary structure (Zuker 2003; Reuter and Mathews 2010; Lorenz et al. 2011). *PinMol* uses as input the “ss-count” file generated by *mfold* upon folding the RNA target sequence of interest. This file lists for each of the target RNA nucleotides (nt) the number of structures out of the total number predicted (comprising the MFE and suboptimal structures) in which the corresponding nucleotide is single-stranded.

#### User-defined parameters

The user selects (Fig. 2):

1. **Probe length** sequence complementary to RNA target, a value between **18** and **26** nt.
2. **Number of desired molecular beacons** to be designed - between **2** and **50,** or maximum number of probes that meet the design criteria, whichever is smaller.
3. **Target region** (optional) for molecular beacon design, while still considering the predicted global structure. This is useful if there are stretches within the mRNA target that are known to be subject to regulation in the cell or tissue of interest (*e.g.* regions that interact with protein factors), which may render them inaccessible for probe hybridization, and thus should not be taken into account for probe design.
4. **Cross-homology** with other RNA targets (optional) – this analysis is performed using an user-generated, external xml file from NCBI “blastn” application.

**FIGURE 2.**
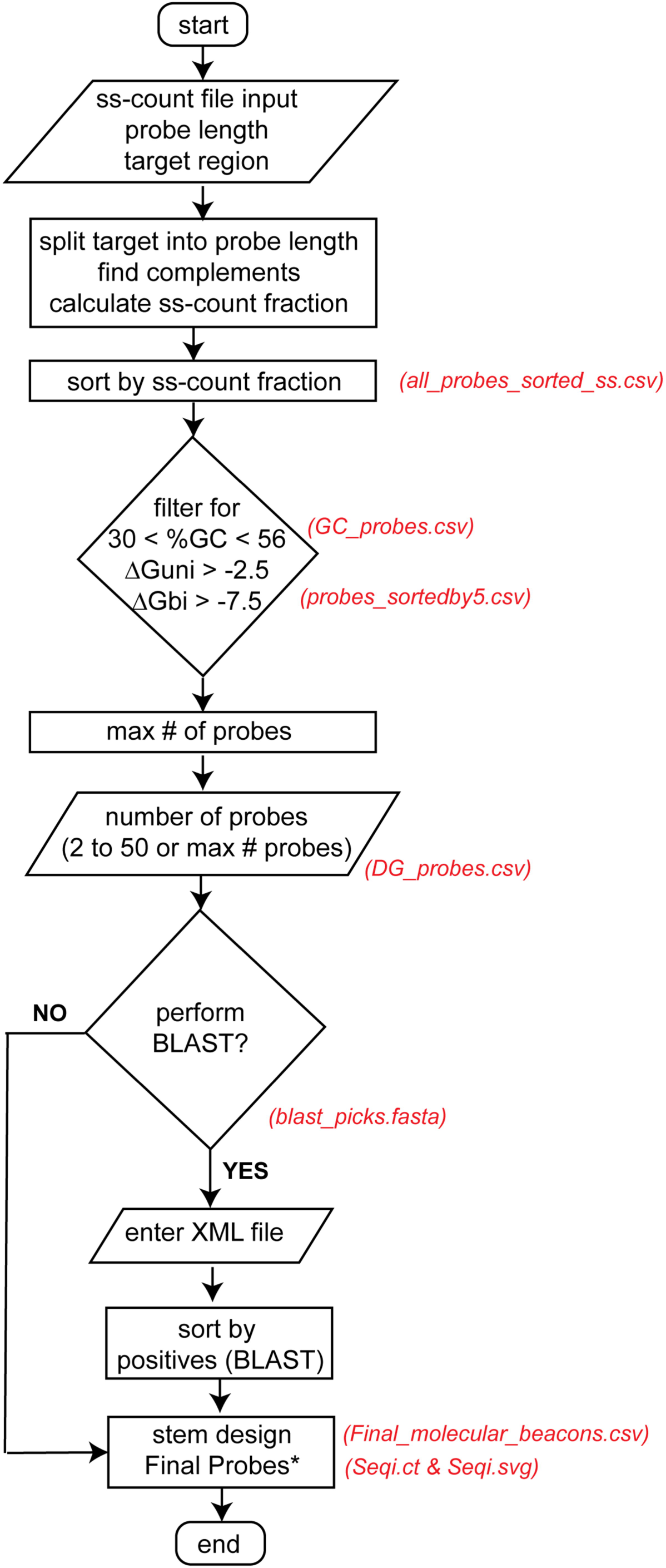
Simplified flow chart for the *PinMol* application. Intermediate files (red) that are generated and saved by *PinMol*. ΔG_uni_ = free energy of unimolecular folding or ΔG_unimol_, ΔG_bi_ = free energy of bimolecular folding or ΔG_bimol_. (*) Molecular beacons that are predicted to fold in a highly structured MFE structure (< −7.2 kcal/mol), or that fail to fold into a hairpin shape, are eliminated. Output files for *oskar*-specific molecular beacons with 24 nt probe length are available for download on our website: https://bratulab.wordpress.com/software/.

#### *PinMol* probe ranking and selection

Initially, all possible probes for the user-selected probe length are sorted according to a target “ss-count fraction”. We define this fraction as the ratio of the sum of the ss-count of each nucleotide [*ss-count(nt*_*i*_)] for a selected probe length to the product of the probe length *(l)* and the number of secondary structures included in the input file (*n*_*str*_) (Eq. 1).

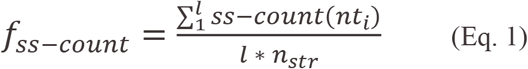

Therefore, *f*_*ss-count*_ is equal to one for a fully single-stranded target region, and zero for a fully double-stranded target region. From these sorted probes, the program filters out the probes with a percentage of GC nucleotides smaller than 31, or greater than 55 (Bratu et al. 2011). For the probes that meet these requirements, *PinMol* calculates the free energy of unimolecular (ΔG_unimol_) and bimolecular (ΔG_bimol_) folding using the *oligoscreen* subroutine of the *RNAstructure* text interface application (Mathews 2006; Reuter and Mathews 2010). Molecular beacons used in live cell imaging experiments are often synthesized with a 2’-*O*-methyl RNA (2’OMe) modified backbone to ensure resistance to cellular nucleases. Therefore, we used RNA parameters to calculate folding free energies, as they more closely match 2’OMe/RNA hybridization profiles when compared to DNA parameters (Tsourkas et al. 2002). To find probes with maximum efficiency for target detection, *PinMol* filters out probes that have a ΔG_unimol_ < −2.5 kcal/mol and ΔG_bimol_ < −7.5 kcal/mol, in order to minimize probe-target hybridization constraints that might be imposed by a highly structured probe region (Fig. 2).

#### Stem design

After the probes have been selected, the stem is designed to form five G/C-rich base pairs. The program takes into consideration if any of the end nucleotides of the selected probe sequence are complementary to each other (up to three on each end), in which case the stem is adjusted to incorporate them. This means that when such complementarity is found, the probe region that is single-stranded may be shorter than the originally selected probe length.

The *PinMol* stem design function also searches for possible complementarity of the newly added stem sequence with the selected probe sequence. When such complementarity is found, the stem sequence is redesigned to ensure that the molecular beacon will fold into a hairpin shape. After the stem is added, each probe is folded and the MFE structure is drawn using the *fold* and *draw* subroutines of *RNAstructure*, respectively (Mathews 2006; Reuter and Mathews 2010). Highly structured molecular beacons are discarded by filtering out the structures with a folding MFE < −7.2 kcal/mol (Fig. 2). In addition, a few molecular beacons that do not fold in the desired hairpin conformation can be obtained, especially for shorter probe lengths. Therefore, after the above described filtering steps, *PinMol* also filters out probes for which the first and the last nucleotides of the molecular beacon sequence are not base-paired in the corresponding “ct” output files generated by the *fold* subroutine.

Although *PinMol* does not include the selection of the fluorophore/quencher pair, when designing molecular beacons it should also be taken into consideration that some pairs have a stabilizing effect (*e.g.* for the same molecular beacon sequence Cy5/BHQ-2 increases the melting temperature (Tm) by 5°C as compared to Cy5/Dabcyl) (Marras et al. 2002).

#### *mfold-* vs *RNAstructure*-generated input files

Three popular RNA folding programs use different thermodynamic parameter sets and therefore, do not predict identical MFE structures, but they are accepted to have similar prediction accuracies. As a consequence, probe ranking using *PinMol* is also expected to differ when using input files generated with each of these programs. However, regardless of the program used to generate the *PinMol* input file, valid molecular beacon candidates will be obtained. We used *mfold* to obtain the input file for *PinMol* because it is the only one that directly generates the ss-count file. The free energies used for filtering were calculated using *RNAstructure,* and thus we also considered target secondary structures predicted using *RNAstructure*. We compared results when using as *PinMol* input the target MFE structure alone (referred to as “MFE”), or both the MFE and the suboptimal secondary structures with 10% of MFE (*e.g.* referred to as “31STR” for *oskar* mRNA) predicted with *mfold* or *RNAstructure*.

#### Defining an optimum number of selected probes

To define an optimum number of probes without limiting the *f*_*ss-count*_, we determined the number of probes that have a *f*_*ss-count*_ > 0.5 for *oskar* (*osk*) mRNA (Fig. 3A), a maternal fruit fly mRNA that we have successfully visualized in live fruit fly egg chambers using molecular beacons (Bratu et al. 2003; Mhlanga et al. 2009; Catrina et al. 2016). We determined the *f*_*ss-count*_ for all *osk* probes with a probe length ranging between 14 and 28 nt using target structural information from four different *PinMol* input files, which were obtained with both *mfold* (MFE, 31STR) or *RNAstructure* (MFE, 31STR). The maximum number of *osk* probes with a *f*_*ss-count*_ > 0.5 was obtained for the 31STR *mfold* input for a probe length of 14 nt (725 vs 706 probes for 31STR *RNAstructure* input). The minimum number of *osk* probes with a *f*_*ss-count*_ > 0.5 was obtained for the 31STR *RNAstructure* input for a probe length of 28 nt (445 vs 511 probes for 31STR *mfold* input). We found that limiting the number of probes to 50, although it may reduce the number of unique probes, ensures a high *f*_*ss-count*_ for target RNAs of an average length (*e.g.* 2,200 nt for mammalian mRNAs). For shorter RNA targets, we recommend that the *f*_*ss-count*_ corresponding to the regions targeted by the selected molecular beacons be evaluated on a case by case basis (Fig. 2: “probes_sortedby5.csv” output file).

**FIGURE 3.**
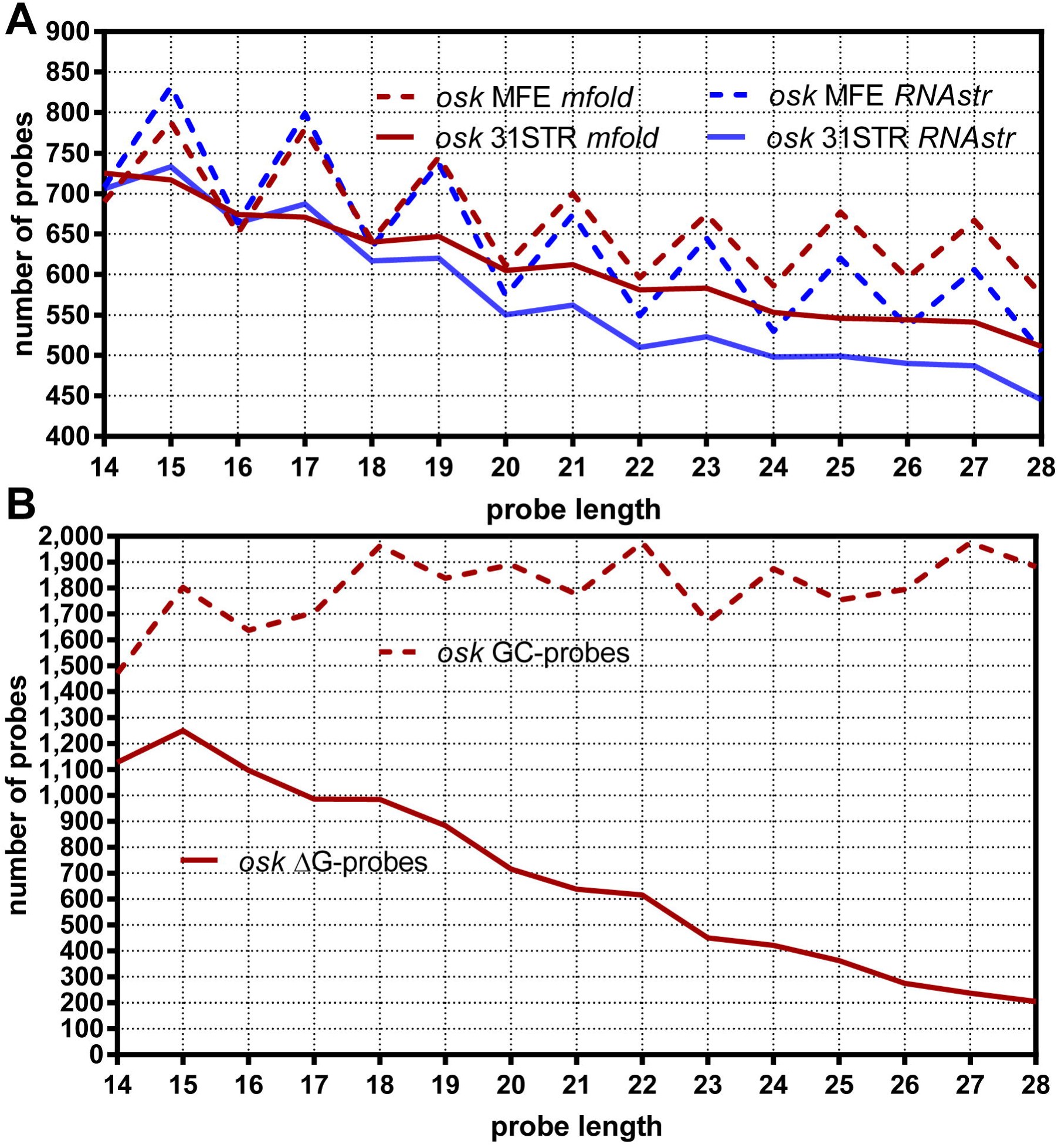
The number of probes generated using *PinMol* for *osk* mRNA target (2,869 nt). (A) The number of probes of a given length (between 14 and 28 nt) that target a region with a *f*_*ss-count*_ larger than 0.5 (more than half of the bases are single-stranded) when using *osk* mRNA *mfold* predicted secondary structures (MFE *mfold –* red dashed, and 31STR *mfold –* red solid line), or *osk* mRNA *RNAstructure* predicted secondary structures (MFE *RNAstr –* blue dashed and 31STR *RNAstr –* blue solid line). (B) The number of probes, for a given probe length, that meet the GC-content requirement (between 31 and 55%) (GC-probes *–* red dashed line), and number of probes that meet the GC-content requirement and the ΔG_unimol_ and ΔG_bimol_ limits (< −2.5 and < −7.5 kcal/mol, respectively, ΔG-probes *–* red solid line) for *osk* mRNA.

Next, we determined the number of probes for *osk* mRNA that meet the GC-content (GC-probes), or both the GC-content and the folding energy (ΔG-probes) requirements, as described above, for probe lengths between 14 and 28 nt (Fig. 3B), and we found that more than 50 probes meet all the design criteria, for each probe length. Since probes are not selected or eliminated based on their *f*_*ss-count*_, the number of probes that meet the GC-content requirement, or meet the GC-content and energetic requirements was independent of the number of structures taken into consideration in the input file (MFE or 31STR) used to run *PinMol* (Fig. 3A vs 3B).

### Cross homology with other RNAs

In microarray assays using 50 nt-long oligonucleotides, the maximum allowed cross complementarity in order to avoid false positive signals is 16 consecutive nucleotides (He et al. 2005). From studies with mismatched molecular beacons, it was shown that no hybrids (molecular beacon/target) are formed even with one nucleotide mismatch, usually placed in the center of the probe sequence (Tyagi et al. 1998; Chen et al. 2007). When considering that the maximum probe length allowed by *PinMol* is 26 nt, such overlap should be less than 13 consecutive nucleotides for molecular beacons of maximum probe length, and this requirement can be relaxed for molecular beacons with probe lengths shorter than 26 nt. Additional information, specific to the model system of interest, should also be taken into consideration. For example, when imaging transcripts expressed in the fruit fly ovary, any homology overlap with other mRNAs not expressed in this tissue can be disregarded. Although *PinMol* includes an optional ranking of the chosen probes using the “xml” output file from a BLAST alignment search, it does not take into account the cell/tissue expression levels of the target. The probe ranking using BLAST alignment results is performed after the desired number of probes are selected, thus these probes are re-sorted in ascending order of the maximum number of positives found for a complementary match between each of the selected probe sequence and an mRNA other than the target of interest. Therefore, this feature is convenient when designing probes targeting ubiquitously expressed mRNAs, but may not be needed if the target mRNA has tissue-restricted expression levels. *PinMol* generates a file containing a list of the selected number of probes in the “FASTA” format, which can be uploaded in the NCBI BLAST form. For mRNA targets, “blastn” search is recommended, using the reference RNA sequences or refseq_rna database for the organism of interest and default parameters. For other RNA targets, the “blastn” search with the nucleotide collection (nr/nt) and default parameters should be used, as the program will prematurely terminate if it does not find at least one fully complementary match to each probe sequence.

*PinMol* provides ranks of the selected number of probes before and after BLAST alignment. Therefore, if BLAST is performed for the chosen probes, it is recommended that the target’s tissue expression level for the corresponding organism is also taken into consideration, and that probes that were ranked based on cross homology results are carefully compared with the pre-BLAST ranking results. This comparison will identify candidate probes that ranked highly before BLAST, but which were placed lower on the list after BLAST, due to a cross homology with an mRNA that is not expressed in the tissue of interest.

### Comparison with experimental data

Molecular beacons have been demonstrated to be highly specific probes, which are able to discriminate against a single nucleotide mismatch between a probe and a target region (Tyagi et al. 1998; Bratu et al. 2003). A false positive signal resulting from a probe hybridizing to a different target is likely to occur in conditions where such mismatches are tolerated. These conditions are minimized in classical RNA-FISH (fluorescence *in situ* hybridization) experiments by using denaturing reagents and increasing the experimental temperature. However, for shorter probes, such as molecular beacons (18-26 nt), the penalty for mismatches greatly increases, and when combined with the energy requirement to open the stem of a hairpin, it allows for specific hybridization at physiological conditions, with target accessibility as the limiting factor.

#### Molecular beacons specific to *osk* mRNA

We have successfully visualized and analyzed *osk* mRNA transport and localization within the *Drosophila melanogaster* egg chamber using fluorescence confocal microscopy coupled with molecular beacons (Mhlanga et al. 2009). The *D. melanogaster* egg chamber goes through 14 stages of development, from germarium to stage 14, and it is composed of somatic (follicle) and germline (one oocyte and 15 nurse cells) cells (McLaughlin and Bratu 2015). Several mRNAs have been reported to asymmetrically localize in the oocyte during oogenesis, and these processes are essential for the development and survival of the future embryo (Lasko 2012). *osk* mRNA is responsible for germplasm determination (Ephrussi et al. 1991), and it is one of the most studied mRNAs in fruit flies. It is actively transported from the nurse cells, and localized and anchored at the oocyte’s posterior during mid- and late oogenesis (stages 7-9) (Ephrussi et al. 1991; Kim-Ha et al. 1991). Therefore, we selected *osk* mRNA as our target mRNA to compare the ranking of molecular beacons designed using *PinMol* with our previously manually designed molecular beacons (Bratu et al. 2003), using both *in vitro* and *in vivo* experiments.

After design, synthesis and purification, the performance of a molecular beacon that is to be used in live cell imaging experiments is characterized by analyzing its 1) thermal denaturation profile and 2) hybridization reaction with a complementary DNA oligonucleotide target and/or *in vitro* transcribed mRNA target. The thermal denaturation indicates whether the molecular beacon is properly folded into a hairpin shape, while the hybridization analysis estimates the binding efficiency of the probe to its target, as well as the fluorescence signal-to-background ratio (Bratu et al. 2011). The efficiency of molecular beacons that were manually designed to target several distinct regions within *osk* mRNA was determined using *in vitro* hybridization reactions with *in vitro* transcribed and folded full-length *osk* mRNA [Table 2, (Bratu et al. 2003)].

**TABLE 2.**
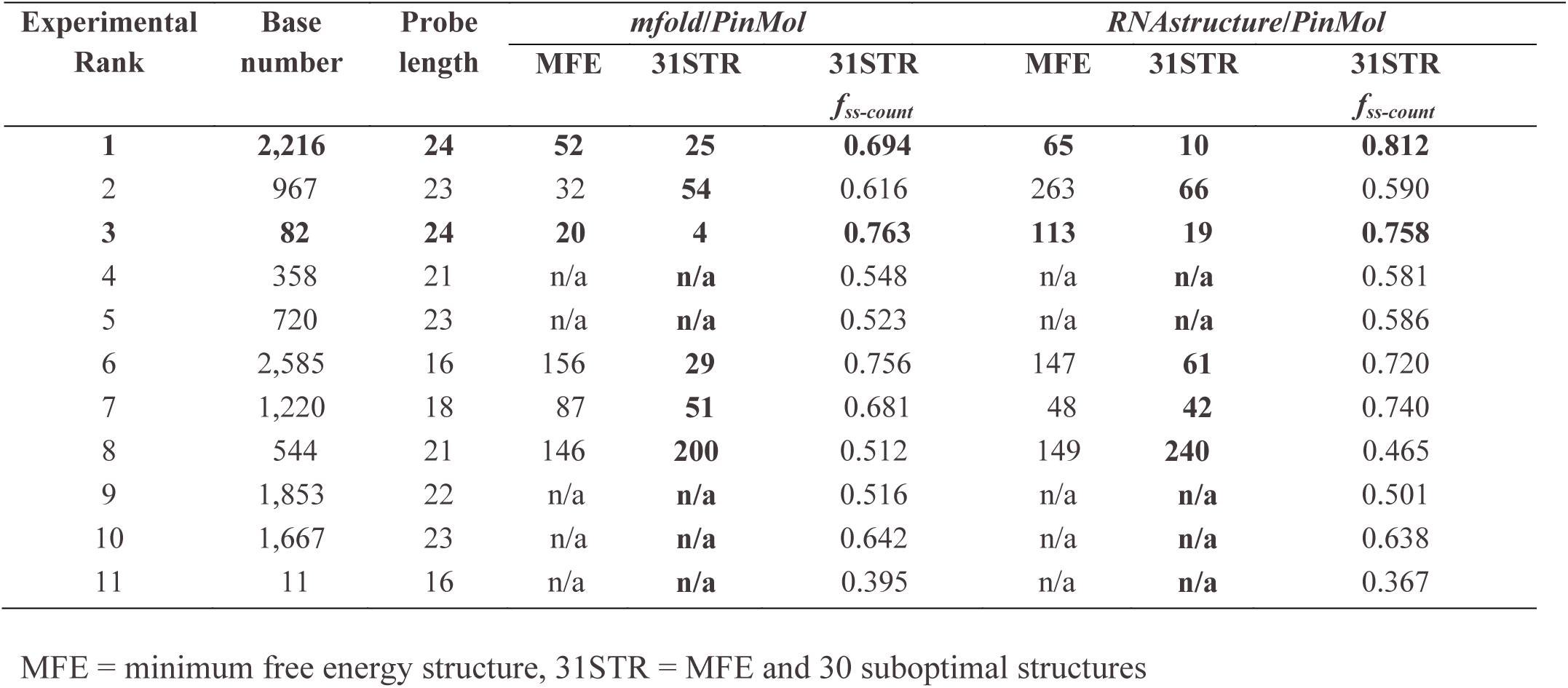
Ranking of *osk* mRNA specific molecular beacons according with their performance in *in vitro* hybridization experiments using full-length *osk* RNA.

| Experimental Rank | Base number | Probe length | *mfold/PinMol* |  |  | *RNAstructure/PinMol* |  |  |
| --- | --- | --- | --- | --- | --- | --- | --- | --- |
|  |  |  | MFE | 31STR | 31STR<br>*f<sub>ss-count</sub>* | MFE | 31STR | 31STR<br>*f<sub>ss-count</sub>* |
| **1** | **2,216** | **24** | **52** | **25** | **0.694** | **65** | **10** | **0.812** |
| 2 | 967 | 23 | 32 | **54** | 0.616 | 263 | **66** | 0.590 |
| **3** | **82** | **24** | **20** | **4** | **0.763** | **113** | **19** | **0.758** |
| 4 | 358 | 21 | n/a | **n/a** | 0.548 | n/a | **n/a** | 0.581 |
| 5 | 720 | 23 | n/a | **n/a** | 0.523 | n/a | **n/a** | 0.586 |
| 6 | 2,585 | 16 | 156 | **29** | 0.756 | 147 | **61** | 0.720 |
| 7 | 1,220 | 18 | 87 | **51** | 0.681 | 48 | **42** | 0.740 |
| 8 | 544 | 21 | 146 | **200** | 0.512 | 149 | **240** | 0.465 |
| 9 | 1,853 | 22 | n/a | **n/a** | 0.516 | n/a | **n/a** | 0.501 |
| 10 | 1,667 | 23 | n/a | **n/a** | 0.642 | n/a | **n/a** | 0.638 |
| 11 | 11 | 16 | n/a | **n/a** | 0.395 | n/a | **n/a** | 0.367 |

MFE = minimum free energy structure, 31STR = MFE and 30 suboptimal structures

We compared the experimental ranking of these *osk*-specific molecular beacons with the ranking of *PinMol* designed molecular beacons for target structures predicted using *mfold* (Supplemental Table S1) or *RNAstructure*, and when considering MFE or 31STR for *osk* mRNA. The ranking of the probes designed for 31STR were the closest to the experimental results, followed by MFE for both programs (Fig. 4). Several probes were chosen for the 1,800-1,900 nt region for MFE *mfold* and *RNAstructure* and 31STR *RNAstructure*, which were found to perform poorly experimentally (Fig. 4). However, when using 31STR *mfold*, no probes were selected for this region (Fig. 4). We found minor differences in the distribution of the top 50 probes when using *mfold-* vs *RNAstructure-*generated input files (Fig. 4), observation consistent with the fact that different folding packages use different sets of RNA parameters, but still lead to similar accuracies in secondary structure predictions. Specifically, certain regions were favored by *mfold* (500-600 nt) and others by *RNAstructure* (1,000-1,200 nt) (Fig. 4-5A).

**FIGURE 4.**
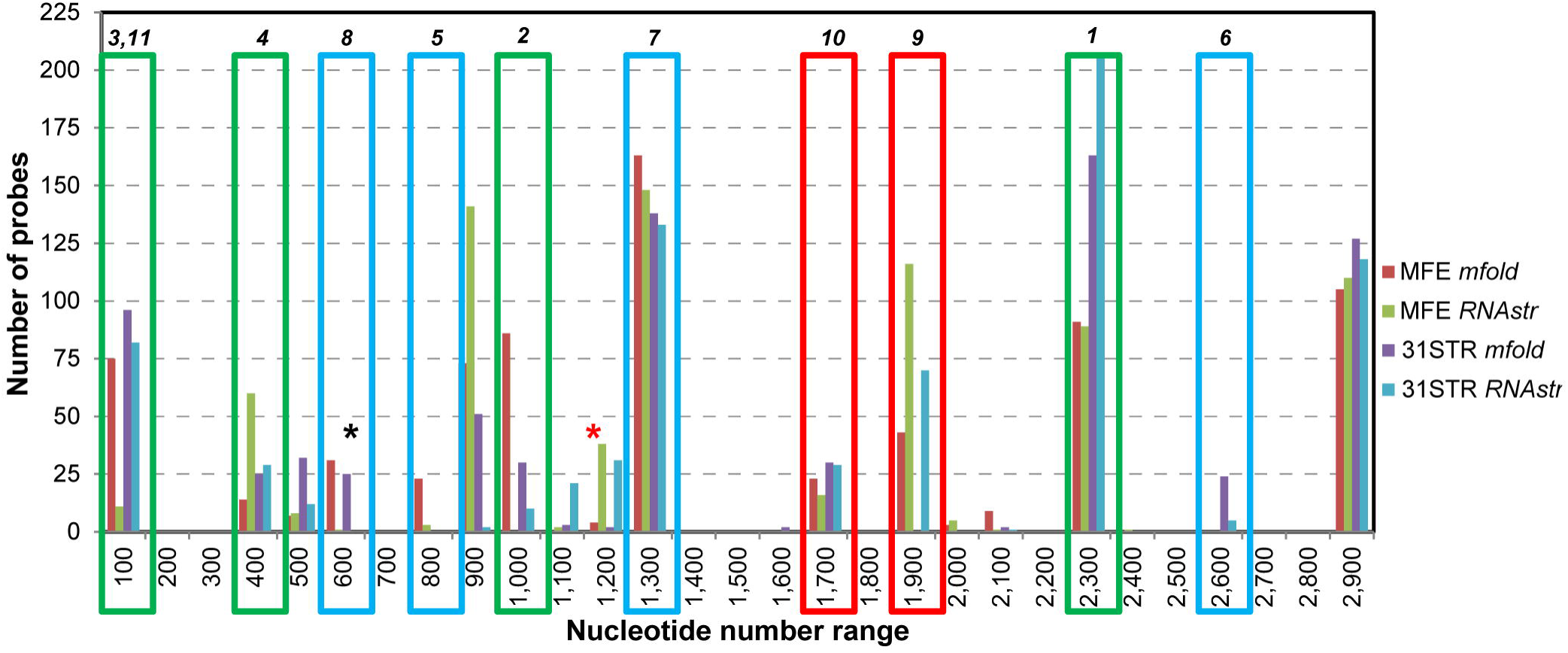
Comparison of target regions selected by the *PinMol* program for designing *osk* mRNA molecular beacons (bars) with experimental results (boxes). (A) The distribution of the probes (bars) designed using a target ss-count file generated with *mfold* (MFE *mfold –* red, 31STR *mfold –* purple,) or *RNAstructure* (MFE *RNAstr –* green, 31STR *RNAstr –* blue,), for probe lengths between 14 and 28 nt. Red boxes show molecular beacons that performed poorly experimentally, blue boxes show probes that have a moderate efficiency to hybridize to their target region, and green boxes highlight probes that hybridize to their target region with the highest efficiency. The numbers on top of the boxes are the *in vitro* experimental (RNA hybridization) rankings of each probe, with one being the most efficient probe. Red and black asterisks highlight nucleotide ranges where more probes were designed using the *RNAstructure* or *mfold* ss-count files, respectively, when considering 30SO structures in addition to the MFE structure.

We also compared the top ten 24 nt-long probe sequences designed when using MFE or 31STR input files with previous experimental results, and found that for 31STR *RNAstructure* all probes were mapped to the same region of the probe experimentally ranked as #1 (osk2216), but probes selected for 31STR *mfold* were from four different regions (100s, 1,200s, 2,200s and 2,800s) (Fig. 5B). Overall, our analysis suggests that the ranking of *PinMol* designed molecular beacons was better aligned with experimental data when 30SO structures were considered in addition to the MFE structure (Table 2; Fig. 4). Moreover, the *PinMol* rank of the top experimental probe improved when using 31STR vs MFE (Table 2; osk2216), while probes with poor experimental performance were eliminated during the filtering steps (Table 2; osk544, osk1853 and osk1667).

**FIGURE 5.**
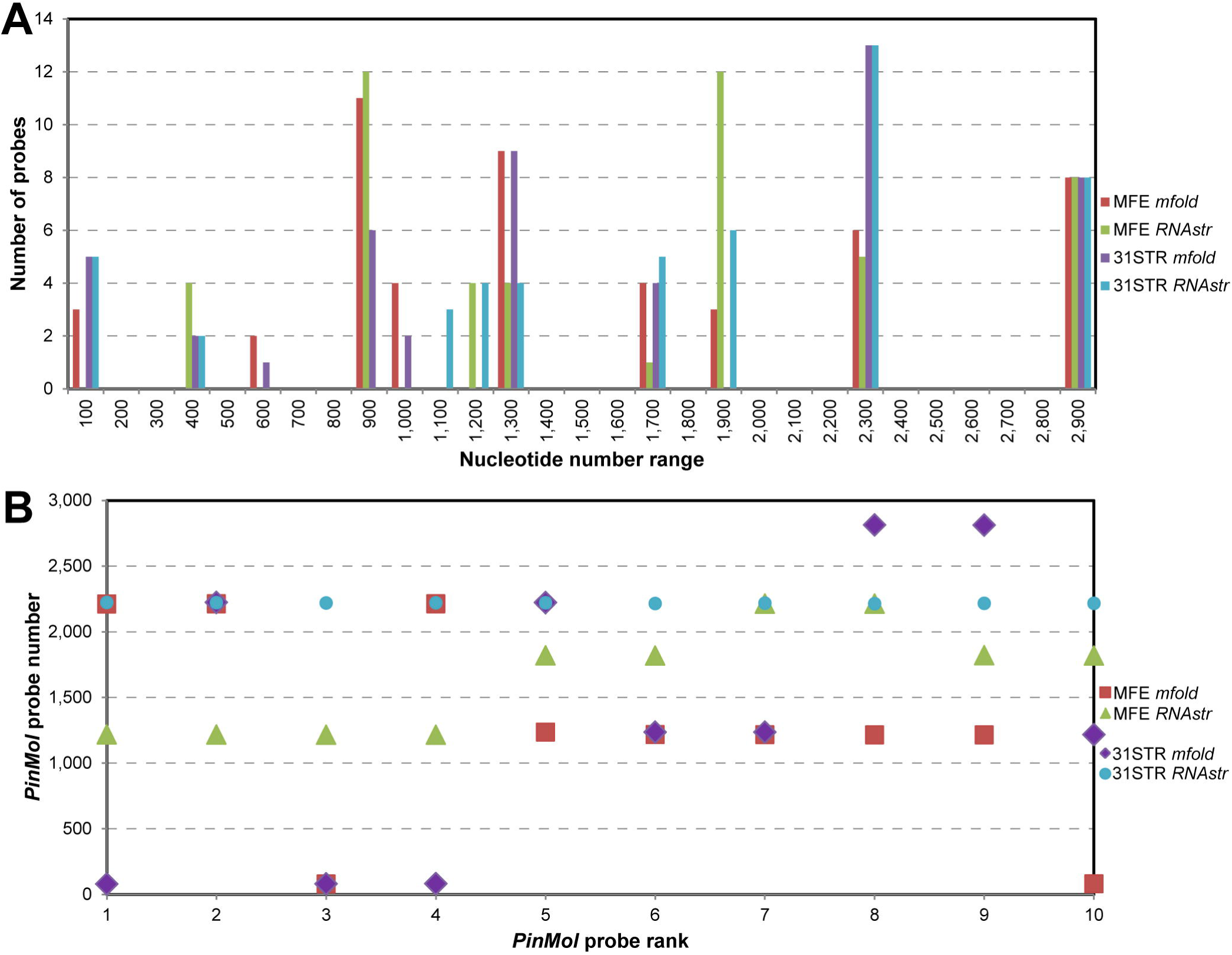
Molecular beacons with 24 nt probe length designed with the *PinMol* program for *osk* mRNA. (A) The distribution of the probes (bars) designed using a target ss-count file generated with *mfold* (MFE *mfold –* red, 31STR *mfold –* purple) or *RNAstructure* (MFE *RNAstr –* green, 31STR *RNAstr –* blue). (B) Top ten probes designed with *PinMol* for the input files indicated in (A).

For detection of low abundance targets, it may be necessary to increase the number of fluorophores per RNA molecule by using multiple molecular beacons that bind to different regions along the target RNA. We found that the 50 designed probes were almost entirely encompassed within ten, strongly supported experimentally, and predicted to be accessible regions (each 100 nt long) of *osk* mRNA (Fig. 4). This indicates that our program selected some “redundant” probes that showed differences of only a few nucleotides (*e.g.* osk2216 and osk2215). While it may be tempting to eliminate these redundant probes, we advise that this should not be done prior to performing BLAST alignment for determining cross homology with other mRNAs. Moreover, it should be considered that the *f*_*ss-count*_ decreases for probes ranked low in the initially filtered probe list (Fig. 2; “DG_probes.csv” *PinMol* output file). Although the stem design is limited to 50 total probes, *PinMol* saves the intermediate files containing all sorted probes after filtering (Fig. 2; “probes_sortedby5.cvs”), which can be used to further inspect the results, and if desired, to select smaller target regions, and then to re-design the probes.

As an example, we selected the region between 1,301 and 1,600 nt of *osk* mRNA, for which 24 nt-long probes were not selected in the top 50 probes designed using *PinMol* (Fig. 5A). For this region, when using 31STR *mfold* as input file the total number of possible probes that are 24 nt-long was 278, out of which 231 met the GC-content requirement, and of which 36 also met the folding energy requirements, but only 15 probes had a *f*_*ss-count*_ > 0.5. The top choice before applying any filters was at base number 1,381 with a *f*_*ss-count*_ of only 0.547, which was ranked 324^th^ in the list of probes sorted in descending order by *f*_*ss-count*_, but osk1381 was eliminated during the initial filtering steps. The final top choice for this region was the probe at base number 1,308, with a *f*_*ss-count*_ of 0.520, and it was ranked 448^th^ when considering the probes selected by *PinMol* for the full-length target *osk* mRNA. Our analysis showed that even the best molecular beacon that *PinMol* designed for this region targets a stretch that consists of single stranded nucleotides for only ~half of its length. This confirms that the selected target region was predicted to be highly structured, and therefore it is not suitable for targeting *osk* mRNA in both *in vitro* and *in vivo* experiments.

### Optimizing the molecular beacon design for live cell imaging

Ideally, when designing molecular beacons for live cell imaging experiments, the *in vivo* target mRNA secondary structure should be taken into consideration for selecting the probe sequence. While *in vivo* mapping has proved useful in determining mRNA secondary structure in live cells and tissues (Mathews et al. 2004; Rouskin et al. 2014), due to the scarcity of such data, molecular beacon design relies on mRNA secondary structures predicted via free energy minimization approaches. Moreover, it was reported that active unfolding of mRNA structures occurs *in vivo* (Rouskin et al. 2014). For *osk* mRNA probe design, when including structural information from 31STR vs MFE, *PinMol* allowed identification of additional regions that are viable targets in *in vivo* experiments, by taking into consideration the mRNA’s folding flexibility.

To test the ranking accuracy of *PinMol* selected probes when using *mfold*-generated input files, we selected molecular beacon osk1236 with a probe length of 24 nt (for which the 5’ end of the stem coincidently includes three additional nucleotides matching the target) to compare it to osk82 and osk2216 (Fig. 6A). We chose the probe length of 24 nt in order to be consistent with the probe length for both, osk82 and osk1236. The osk2216 molecular beacon had the best performance, while osk82 placed third in *in vitro* hybridization experiments (Table 2) (Bratu et al. 2003), and osk2216 was also efficient in detecting endogenous *osk* mRNA in *in vivo* experiments (Catrina et al. 2012). When comparing their *f*_*ss-count*_, osk1236 was predicted to outrank osk2216 in all cases with the *mfold* input files, but the opposite trend was obtained with the *RNAstructure* input files (Table 3). However, osk82 outranked osk1236 when using the *mfold* 31STR ss-count file. When using *RNAstructure*-derived input files (MFE and 31STR) osk2216 outranked both osk82 and osk1236, consistent with the experimental *in vitro* RNA hybridization results (Table 2).

**FIGURE 6.**
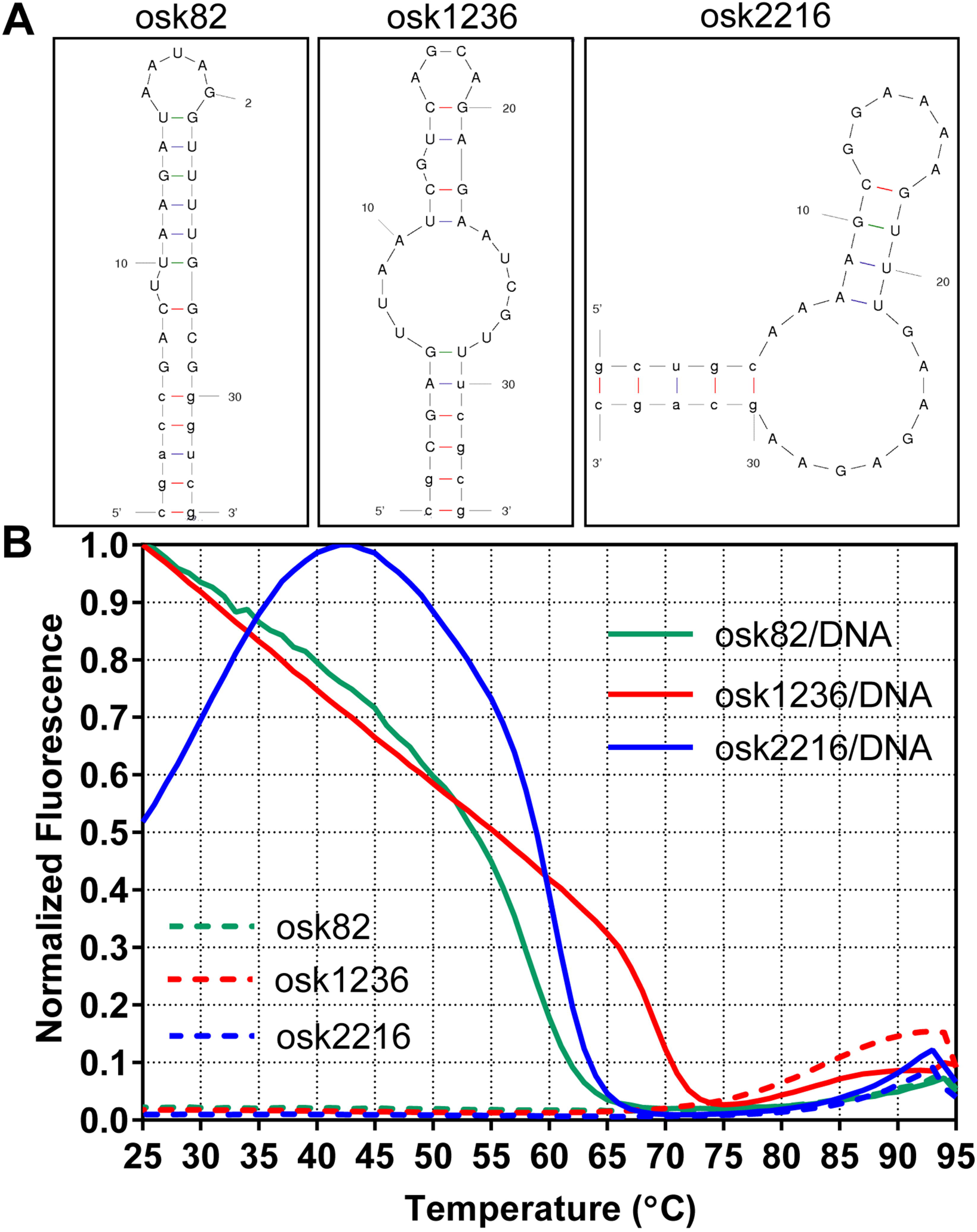
Comparison of the thermal denaturation profiles of molecular beacons osk1236, osk82 and osk2216. (A) The *mfold* predicted MFE structure for each molecular beacon. (B) Thermal denaturation curves for molecular beacon alone (dashed lines), or molecular beacon and DNA oligonucleotide target (solid lines): osk82 (green), osk1236 (red) and osk2216 (blue).

**TABLE 3.** Comparison of parameters for the probe sequence of three molecular beacons targeting different regions of *osk* mRNA.

| Parameters | MFE |  |  | 31STR |  |  |
| --- | --- | --- | --- | --- | --- | --- |
|  | osk82 | osk1236 | osk2216 | osk82 | osk1236 | osk2216 |
| ***mfold*** |  |  |  |  |  |  |
| $f_{ss-count}$ | 0.750 | 0.792 | 0.667 | 0.763 | 0.743 | 0.694 |
| Rank | 20 | 5 | 52 | 4 | 6 | 25 |
| ***RNAstructure*** |  |  |  |  |  |  |
| $f_{ss-count}$ | 0.542 | 0.500 | 0.667 | 0.758 | 0.504 | 0.812 |
| Rank | 113 | 125 | 65 | 19 | 103 | 10 |

MFE = minimum free energy structure, 31STR = MFE and 30 suboptimal structures

We synthesized and purified osk1236 (MATERIALS AND METHODS) and when comparing the thermal denaturation profiles for these three molecular beacons, we found that osk1236 had a slightly lower Tm than osk2216 and osk82, consistent with *mfold* results (82°C vs 88°C and 86°C, Table 4, Fig. 6B). For thermal denaturation profiles of molecular beacon in the presence of DNA oligonucleotide target, there was a similar trend with osk1236/DNA having a lower Tm than osk2216/DNA, but not osk82/DNA (55°C vs 59°C and 53°C, Table 4, Fig. 6B), although osk1236/DNA contains a higher percentage of G/C base pairs when compared to osk2216/DNA (GC%: osk82, 37.5, osk1236, 41.7 and osk2216, 33.3). The thermal denaturation profile of osk2216 with its DNA oligonucleotide complement is complicated by the combination of the probes’s low percentage of GC nucleotides and moderately favorable ΔG_bimol_ (−17.4 kcal/mol for the osk2216 molecular beacon sequence and −4.4 kcal/mol for the probe sequence). When using a DNA oligonucleotide target, the intermolecular interaction of osk2216 molecular beacon is favored over the formation of osk2216/DNA hybrid, and as a result the molecular beacon falls off the target DNA at temperatures lower than 43°C, which led to a sharp decrease in fluorescence signal (Fig. 6B). Because RNA/RNA hybrids are energetically more stable than RNA/DNA hybrids, this behavior is less likely to occur when the molecular beacon hybridizes with its intended mRNA target region. We found that the hybridization half-lives of osk1236 with the corresponding DNA oligonucleotide target is slightly longer than that of osk2216 and osk82 molecular beacon (12.6 vs 1.2 and 2.6 sec) (Table 4; Fig. 7A). However, for hybridization reactions with full-length, *in vitro* transcribed *osk* RNA, we observed that osk2216/RNA (47.6 sec) and osk82/RNA (28.6 sec) hybrids had a much shorter half-life than the osk1236/RNA hybrid (480.4 sec) (Table 4; Fig. 7B).

**TABLE 4.**
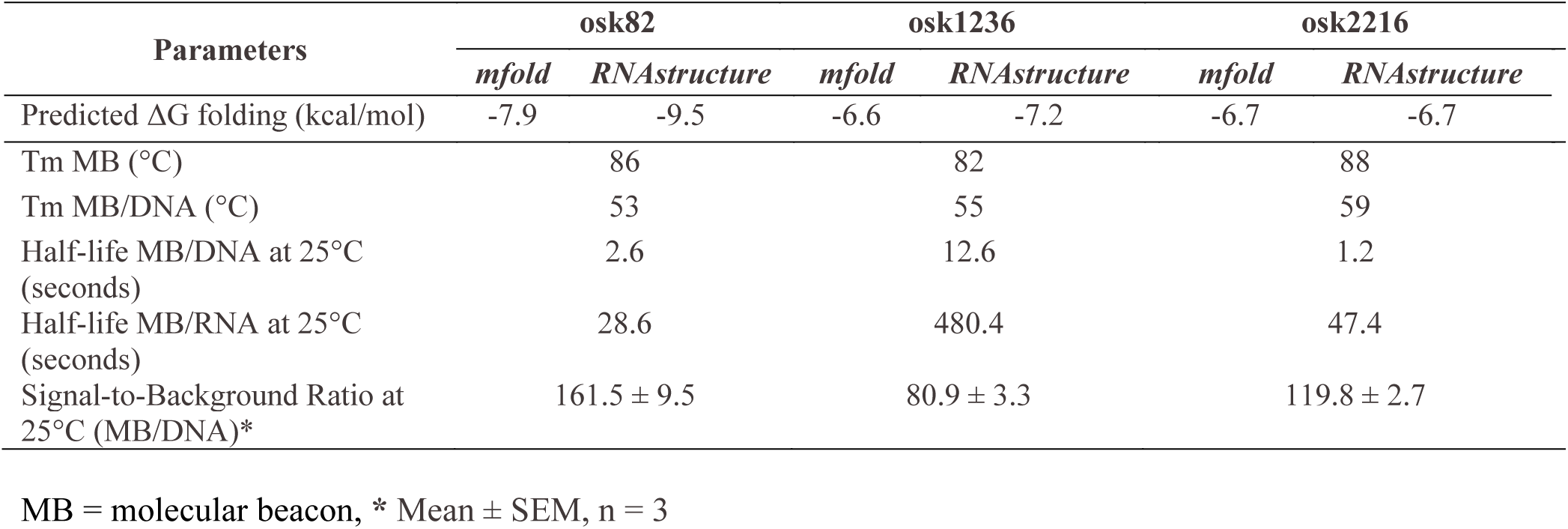
*In vitro* hybridization results for osk82, osk1236 and osk2216 molecular beacons.

**FIGURE 7.**
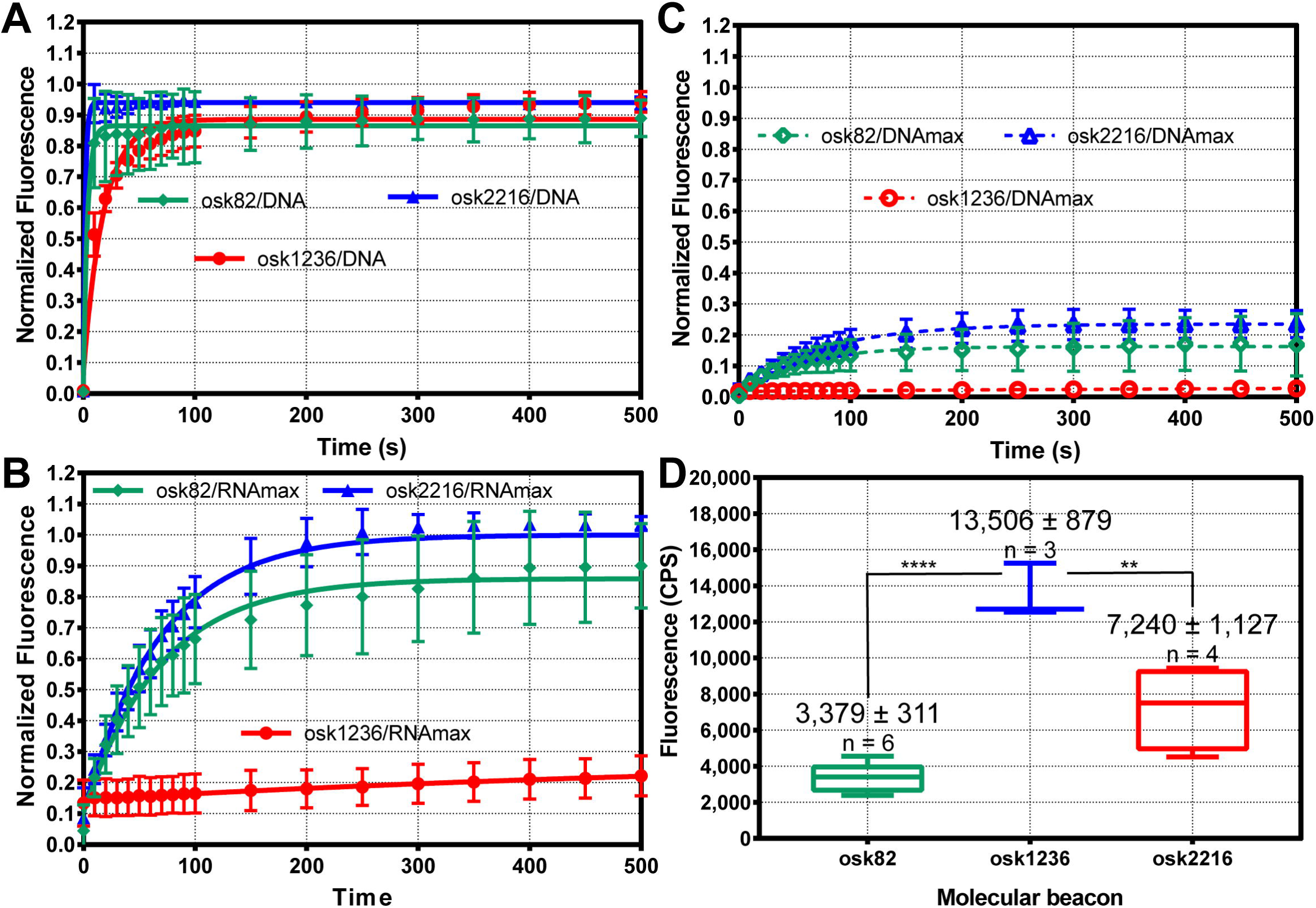
Comparison of *in vitro* molecular beacon/target hybridization profiles of osk1236,s osk82 and osk2216. Hybridization curves for each molecular beacon with the corresponding complementary (A) DNA oligonucleotide target (MB/DNA) (osk82 – green solid line, osk2216 – blue solid line, osk1236 – red solid line, filled symbols), or (B) and (C) with full-length, *in vitro* transcribed *osk* RNA, (osk82 – green, osk2216 – blue, osk1236 – red). In (B) are shown the RNA hybridization curves normalized using the fluorescence signal from first plateau of the RNA hybridization reaction (MB/RNAmax, solid lines, filled symbols). In (C), for comparison, the fluorescence signal (CPS) for the RNA-hybridization curves was normalized to the maximum fluorescence signal obtained upon adding DNA target after the RNA hybridization reaction reached the first plateau (dashed lines, open symbols). (D) Average maximum fluorescence signal (CPS) from *in vitro* RNA hybridization experiments for each molecular beacon (floating bars: osk82, green, osk1236, red, osk2216, blue, mean ± SEM unpaired t-test: ****, p < 0.0001 and **, p < 0.0092). MB = molecular beacon (*i.e.* osk82, osk1236 or osk2216)

We analyzed and quantified the *in vivo* performance of each molecular beacon, osk82, osk1236 and osk2216. Quantification of molecular beacon *in vivo* target detection efficiency is not trivial due to several variables that need to be controlled that are expected to result in a wide variation in the final results, which we observed for osk1236 (Table 5). We took into account some of these variables:

1. **Molecular beacon quantity delivered** – to control for how much molecular beacon is injected in each egg chamber, we added to the molecular beacon injection solution, DNA oligonucleotides labeled with a spectrally different fluorophore (TMR), which we designed for use in RNA single molecule *in situ* hybridization (smFISH) specific to *gurken* mRNA (Bayer et al. 2015).
2. **Proximity of the injection site to the cellular location visualized** – as much as possible the injections were performed within a nurse cell proximal to the oocyte.
3. **Stage of the development of the microinjected egg chamber** – we selected stage 8 to late stage 9 egg chambers for microinjections, in which the *oskar* mRNA target is mainly localized at the posterior end of the oocyte; to account for the variations in the size of the oocyte, we used a correction factor to adjust the calculated relative detection efficiency of a molecular beacon in larger vs smaller oocytes (Eq. 2)
4. **Depth of image acquisition to collect the optical slice with maximum fluorescence signal** – we acquired 14 optical slices (0.3 μm each) and chose the XY-plane with maximum signal in the selected region of interest (ROI).
5. **Timing of the initiation of acquisition protocol** – we initiated acquisition immediately after acquiring the Z-stack for the TMR channel injection control (also see MATERIALS AND METHODS); to control for any delay in the start of data acquisition, we quantified the change in the Cy5 signal intensity within the posterior ROI from time point t1 (0 min) to t3 (4 min).

**TABLE 5.** Relative detection efficiencies, quantification and statistical analysis of molecular beacon *in vivo* imaging results.

| Molecular Beacon | Efficiency ± SEM | $p^a$ | # injections |
| --- | --- | --- | --- |
| osk82 | 0.130 ± 0.015 | *** / 0.0004 | 8 |
| osk1236 | 0.844 ± 0.163 | n/a | 7 |
| osk2216 | 0.162 ± 0.021 | ** / 0.0013 | 7 |

<sup>a</sup> unpaired t-test for comparing *in vivo* detection efficiency of osk1236 with osk82 and osk2216

Despite osk1236’s less efficient *in vitro* performance with target RNA when compared to that of osk2216 and osk82, osk1236 presented a superior fluorescence signal in *in vivo* experiments (Fig. 8; Table 5: 0.84 vs 0.16 and 0.13 relative detection efficiency; Supplemental Movies S1-S4). Although, we used the same fluorophore for all molecular beacons (Cy5), the two previously synthesized molecular beacons (osk82 and osk2216) had an aged fluorophore, which could result in a decreased fluorescence signal as compared to osk1236. To evaluate if this could have impacted our quantitative analysis, we determined how the performance of the three *osk* molecular beacons compare in *in vitro* hybridization experiments with *osk* RNA target. We calculated the average maximum fluorescence for each molecular beacon (Fig. 7D), and determined the expected normalized fluorescence after 4 min, the time used to quantify the *in vivo* fluorescence signal (osk82: 0.8, osk1236: 0.19 and osk2216: 1), which is the fraction of fluorescence signal from maximum possible in the presence of RNA target. After 4min, the maximum raw fluorescence signal (CPS) was obtained for osk2216, followed by osk82 and osk1236 (37% and 35% of osk2216 CPS, respectively). Although based on *in vitro* RNA hybridization results of the three *osk* molecular beacons we would expect osk2216 to yield the highest raw fluorescence signal *in vivo*, we found that osk1236 outperformed both, osk2216 and osk82. This was evident when comparing *in vivo* imaging results for stage 8 egg chambers microinjected with similar amounts of the corresponding molecular beacon solution, which were acquired and processed using the same parameters, and where a bright fluorescence signal is observed only for osk1236 (Supplemental Movie S1).

**FIGURE 8.**
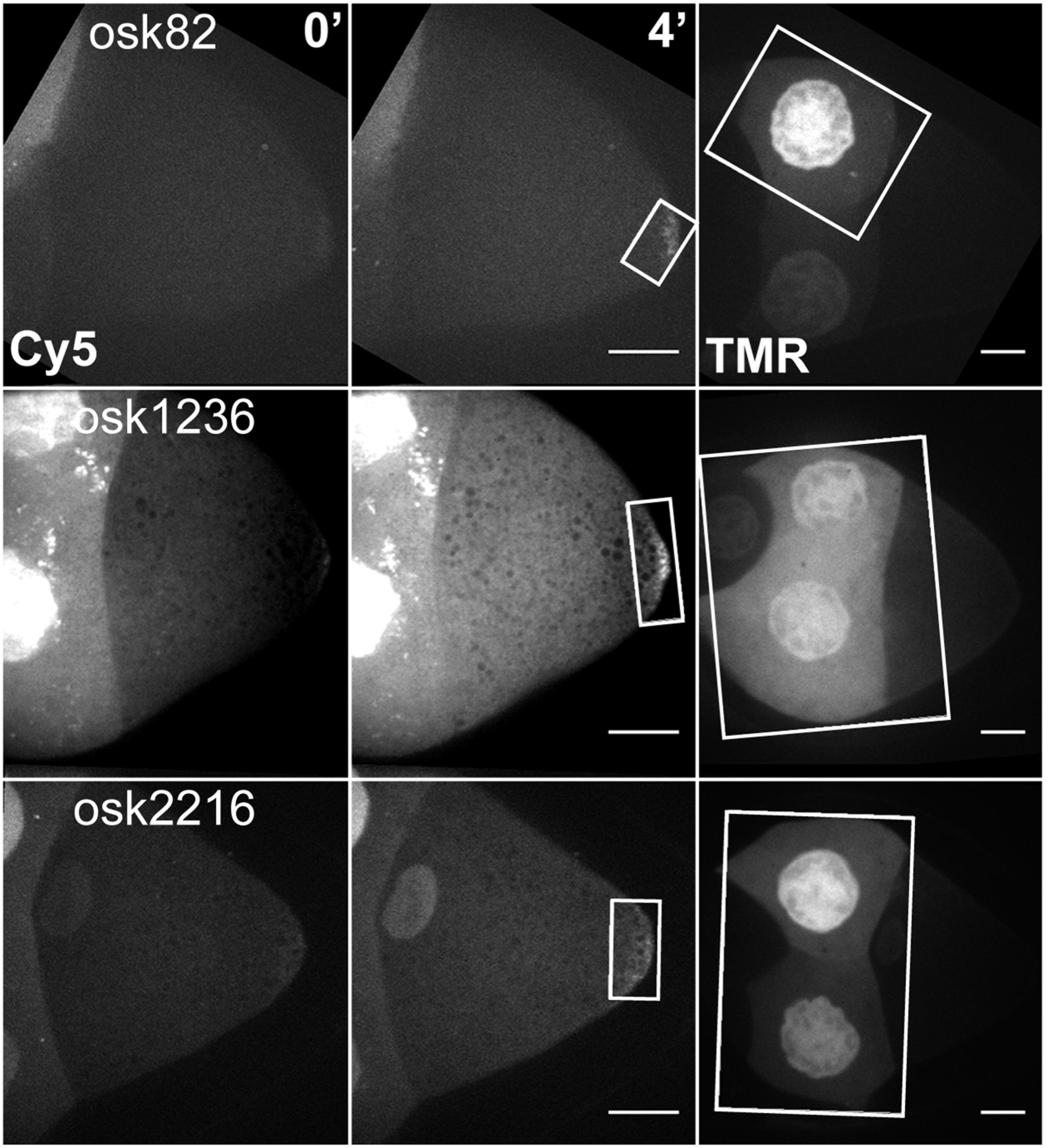
Visualization and quantification of the detection efficiency of endogenous *osk* mRNA in live cell imaging. Fluorescence signal generated by molecular beacon osk82, osk1236 or osk2216 microinjected in late stage 9 wild type fruit fly egg chambers, at the indicated time points. White rectangles indicate the ROIs used for quantification analysis of fluorescence signal generated by each molecular beacon (Cy5), and includes a control for the amount of each injected molecular beacon (TMR). Scale bar is 20 µm.

It is possible that the regions targeted by osk82 and osk2216 are not easily accessible *in vivo* due to protein factors, which would lead to a decreased detection efficiency as compared to *in vitro* results. It is also possible that the RNA region targeted by osk1236 is more rigid in solution than *in vivo*, which leads to a longer osk1236/RNA hybridization half-life, or. Interestingly, *RNAstructure* predicts a lower *f*_*ss-count*_ for osk1236 with 31STR (0.504), or when only considering the MFE structure (0.500) (Table 3). Therefore, when comparing ranking of *PinMol*-designed molecular beacon obtained with *RNAstructure* vs *mfold* input files, osk1236 is ranked only as the 125^th^ vs 5^th^ probe, when using only the MFE, and the 104^th^ vs 6^th^ probe for 31STR. osk2216 is highly ranked by *PinMol* when using *mfold*- or *RNAstructure*-generated input files (25^th^ and 10^th^ probe) when also taking into consideration target suboptimal structures, but it is not selected in the first 50 probes when using MFE as *PinMol* input file (52^nd^ and 65^th^ probe).

Taken together, these results suggest that when incorporating information from predicted target RNA suboptimal secondary structures in addition to the MFE structure, the design of molecular beacon probes for *in vivo* experiments improves (Table 2). Although we recommend using the ss-count file generated by *mfold*, analyzing and comparing target structures predicted by both *mfold* and *RNAstructure* may reveal regions that are consistently predicted to be more accessible, which will maximize the chances of designing highly efficient molecular beacons for live cell imaging experiments.

### Selecting the probe length

*PinMol* allows the design of up to 50 probes with lengths in a range of 18-26 nt. How does one decide what probe length will work best for a particular target or target region? This should not pose a problem when the targeted region is thousands of nucleotides long, but when using a short target, probe selection will be constrained by additional factors. The number of probes meeting all criteria (ΔG-probes) significantly decreased for probe lengths larger than 22 nt (Fig. 3B), indicating that this number may be an optimal length. However, if A/T or G/C rich regions are targeted, in order to meet all requirements as implemented in *PinMol*, a longer or shorter probe length should be selected, respectively.

Ultimately, the main consideration for designing the probe sequence was target accessibility, which was taken into account by sorting all possible probes in descending order of the *f*_*ss-count*_. *PinMol* does not limit the *f*_*ss-count*_ to a lower bound (*e.g.* consider only probes that target a region with a *f*_*ss-count*_ > 0.5) because it limits the total number of selected probes to 50, or the maximum number of possible probes, whichever is smaller. Taking into consideration our previous experimental results, optimization of other parameters (*e.g.* probe length) is also required for the selection of the most efficient probe(s) for live cell imaging. Therefore, it is important to know which factors are most useful in predicting highly efficient probes. We first analyzed our previous experimental data, and we found that only two molecular beacons showed high efficiency in detecting the target region in *in vitro* experiments, while the remaining nine molecular beacons were all under the 50% hybridization efficiency as compared to osk2216, our best experimental probe (Fig. 9) (Bratu et al. 2003). When we compared the top two probes, the *f*_*ss-count*_ appeared to be sufficient to predict their rank (0.694 vs 0.616, 31STR *mfold*) (Table 2). When we compared the first two probes with the third (osk82), although they have similar probe lengths, osk82 had the highest *f*_*ss-count*_ (0.76) when considering 30SO structures in addition to the MFE structure, but it also had a more stable bimolecular folding (Table 2, Fig. 9). This suggests that the ΔG_bimol_ may be another important determinant of probe efficiency for probe lengths in the range of 21-24 nt. The probes ranked experimentally as 4^th^, 5^th^, 9^th^, 10^th^ and 11^th^ were not selected by *PinMol* because they have a ΔG_bimol_ < −7.5 kcal/mol, and the first two probes also have a ΔG_unimol_ < −2.5 kcal/mol. Therefore, these *in vitro* data support our assessment that probes with favorable uni- and/or bimolecular folding will perform poorly (Fig. 9). Of the remaining probes (Table 2), osk2585 ranked 6^th^ and osk1220 ranked 7^th^ showed a decrease in performance that was directly related to their *f*_*ss-count*_, with osk2585 (0.76) having a larger *f*_*ss-count*_ than osk1220 (0.68), and also present a more stable bimolecular folding energy (Table 2; Fig. 9). However, osk2585 showed a slightly lower efficiency than two probes that were filtered out by *PinMol* (osk358 and osk720, experimentally ranked 4^th^ and 5^th^, Table 2). It is possible that the shorter probe length (16 vs 21 and 23 nt) strongly impacted the performance of the former, and thus we chose 18 nt as the minimum length of probe sequence. Finally, the molecular beacon experimentally ranked 8^th^ (osk544) presented the second lowest *f*_*ss-count*_ (0.512), which explains its poor performance in *in vitro* hybridization experiments with full-length *osk* mRNA (Table 2).

**FIGURE 9.**
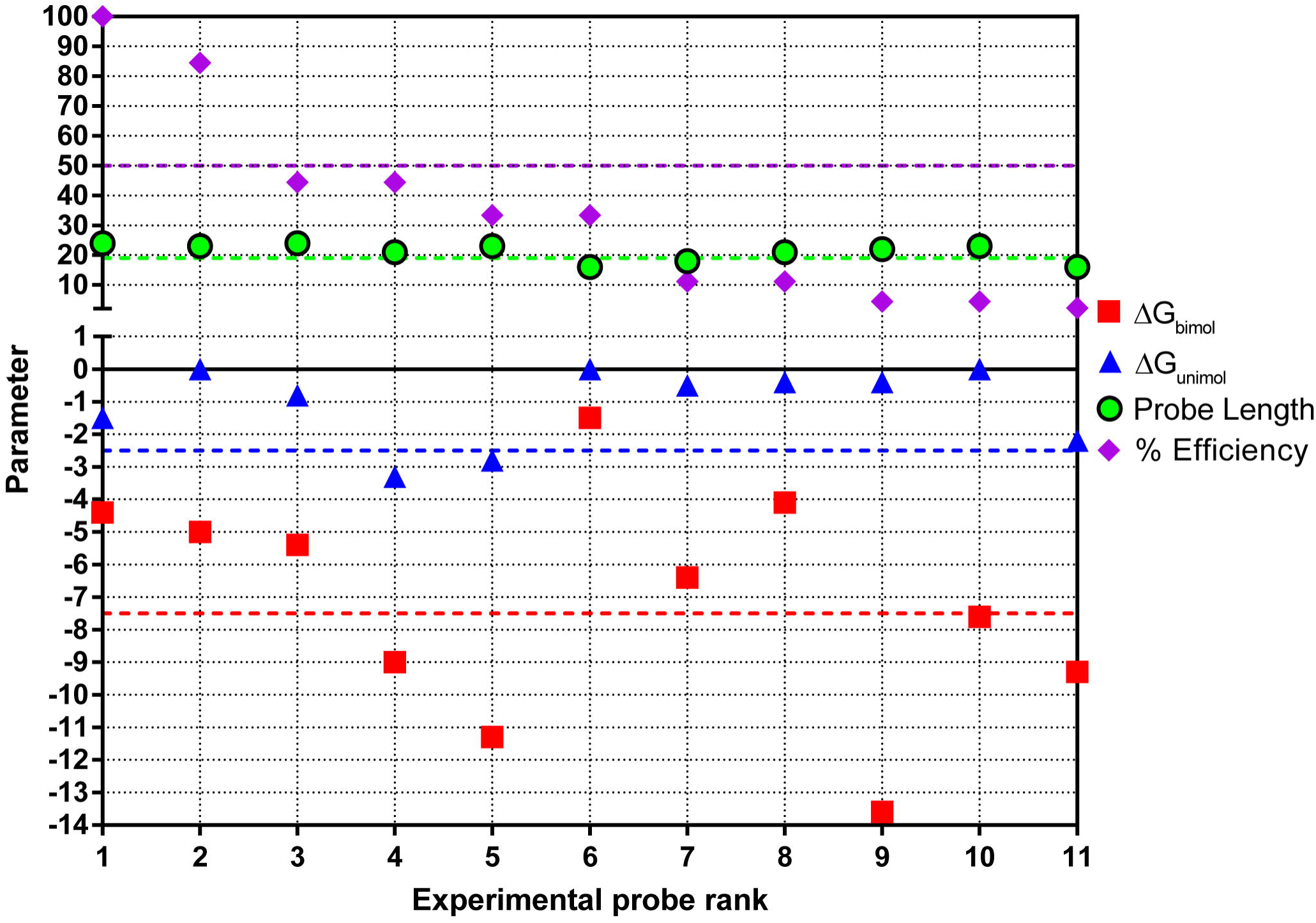
Parameters used by *PinMol* for probe selection. *oligoscreen* predicted values for the probe sequence of experimental molecular beacons previously reported for *osk* mRNA (Table 2) (Bratu et al. 2003). With colored dashed lines are highlighted the limits on probe selection as imposed by *PinMol* for design of molecular beacons suitable for live cell imaging. All probes are shorter than the upper limit of probe length, thus only the lower limit is presented.

Inter- or bimolecular folding is correlated with the molecular beacon’s concentration, and it may negatively impact purification of the probe when using concentrated stock solutions. However, its impact on *in vivo* experiments is expected to be somewhat reduced since lower concentrations are used. Therefore, instead of setting a stringent cut-off for ΔG_bimol_, we chose a moderately low value of −7.5 kcal/mol. For comparison, the ΔG_bimol_ folding of the osk2216 probe region with a fully complementary RNA oligonucleotide was predicted using the *oligoscreen* subroutine to be −35.3 kcal/mol (ΔG_duplex_ in “probes_sortedby5.csv” *PinMol* output file).

We determined the number of molecular beacons designed by *PinMol* for each probe length when using the 31STR input files generated with *mfold* and *RNAstructure*. For all probe lengths, the average final number of molecular beacons was 27 and 30 for *mfold* and *RNAstructure*, respectively.

## DISCUSSION

*PinMol* facilitates the design of molecular beacons for live cell imaging experiments. We have tested the performance of *PinMol* by comparing the ranking of *PinMol* designed molecular beacons with experimental results for manually selected molecular beacons specific for *osk* mRNA, a well-studied fruit fly maternal mRNA. Based on our analysis, we propose the following guidelines when designing molecular beacons for live cell imaging:

1. Use the *mfold*-generated ss-count, and consider target suboptimal secondary structures with a change in folding free energy within 5-10% of the MFE.
2. Avoid probes complementary to the ends of the 5’UTR and 3’UTR of the mRNA target: these regions are usually targeted by cellular factors and/or form long distance base pairs; this is supported by the observed low *in vivo* efficiency for osk82, despite showing a strong performance in *in vitro* experiments.
3. Select the length of the probe sequence between 18 and 26 nt: we propose an optimum length of 22 nt, but this is ultimately dependent on the base composition of the targeted region, which will require longer probes for A/U-rich and shorter probes for G/C-rich sequences.
4. If possible, avoid a G nucleotide at the 5’ end of the molecular beacon: it can quench the fluorescence of certain fluorophores (Marras et al. 2002; Nazarenko et al. 2002).
5. Perform BLAST alignments: carefully evaluate any cross homology by taking into consideration if the target mRNA shows tissue-specific expression.

## MATERIAL AND METHODS

### *PinMol* software

The *PinMol* software was written using Python 3.5.2, for use on Mac and Windows platforms, and packaged for easy download from GitHub (https://github.com/icatrina/PinMol_Mac and https://github.com/icatrina/PinMol_Win). *PinMol* requires Python3, Pandas, Biopython, and text-interface *RNAstructure* (requires free registration and it is available for download from the Mathews lab webpage at University of Rochester: http://rna.urmc.rochester.edu/RNAstructureDownload.html). *PinMol* v1.0 was tagged and released using Git.

A detailed tutorial for the Mac version with or without BLAST analysis, the input files used to design *osk-* and *nanos*-specific molecular beacons with *PinMol*, as well as the output files for designing *osk*-molecular beacons for 24 nt-long probes using *mfold*-generated 31STR input file, are available on our laboratory’s homepage at https://bratulab.wordpress.com/software/.

### Secondary structure prediction

The MFE and suboptimal RNA target secondary structures were predicted using the RNA form of *mfold* with RNA3.5 parameters, or using *RNAstructure* 5.8.1. The ss-count files containing information on MFE and selected suboptimal structures were saved as simple text files. Since there is no ss-count file for the MFE structure alone, we converted the corresponding “.ct” files as follows: we copied the first and second columns and replaced the length of the folded RNA target at the top of the first column with “1”, and added information from column five by replacing the “0” representing a single-stranded nucleotide with “1”, and also replacing a nucleotide number representing a binding partner with “0”. We generated ss-count-formatted files from the *RNAstructure* “.ct” output file (MFE and 31STR) using a Python script, which will be made available upon request.

### Molecular beacon design, synthesis and purification

The molecular beacons osk2216, Cy5-C6-*gcugc*AAAAGCGGAAAAGUUUGAAGAGAA*gcagc*-BHQ-2, and osk82 Cy5-C6-*cgacc*GACUUAAGAUAAUAGGUUUUGGCG*ggucg*-BHQ-2, are the previously described and characterized osk76 and osk2209 molecular beacons (Bratu et al. 2003; Mhlanga et al. 2009). The 2’OMe oligonucleotide labeled with a 5’ C6-amino linker and a 3’ BHQ-2 corresponding to osk1236 sequence, NH2-C6-*cgCGA*GUUAAUCGUCAGCAGAGAAUCGUU*ucgcg*-BHQ-2, was synthesized on the 100 nmole scale by Integrated DNA Technologies. The oligonucleotide was labeled with amino-reactive Cy5 and purified using HPLC as previously described (Bratu et al. 2011). CAPS show the single stranded probe sequence, *italics* show the stem region, and small letters represent the newly added nucleotides as part of the stem region. The sequences for the complementary DNA oligonucleotides used for *in vitro* molecular beacon-target hybridization reaction are as follows: osk82.c CCGCCAAAACCTATCTTAAGTCC, osk1236.c CCTACAACGATTCTCTGCTGACGATTAACTCGGAT, and osk2216.c AAAACTTCTCTTCAAACTTTTCCGCTTTTCCCAA.

### *In vitro* thermal denaturation and hybridization experiments

Thermal denaturation reactions were run with 300 ng (~190 nM) of molecular beacon alone or with 3 µg (~2.3 µM) complementary DNA oligonucleotide target. For molecular beacon thermal denaturation, the reaction was cooled down from 95°C to 25°C, at a rate of 1°C/30 sec. For thermal denaturation experiments in the presence of complementary DNA oligonucleotide target, 3 µl of DNA oligonucleotide target were added to 120 µl containing 300 ng molecular beacon in hybridization buffer (50 mM Tris-HCl - pH 7.5, 1.5 mM MgCl_2_ and 100 mM NaCl). Experiments were performed in a Fluoromax-4 photon counting spectrofluorometer (Horiba-Jobin Yvon), and for each temperature, three time points were acquired every 15 sec, after 0.2 min temperature equilibration with an integration occurring over 0.5 sec. The average of the three readings for each temperature was graphed using GraphPad Prism. The absolute fluorescence intensity (CPS) was normalized for each molecular beacon to the maximum CPS signal obtained for the thermal denaturation of molecular beacon with DNA oligonucleotide target. The excitation and emission slit widths were each 3 nm and 2 nm, respectively, with excitation/emission wavelengths of 649/670 nm.

For *in vitro* hybridization reactions using complementary DNA oligonucleotide targets, 100 ng (~130 nM; 60 µl reaction volume) of the molecular beacon was incubated with 500 ng (~780 nM) of the corresponding complementary DNA oligonucleotide target. The fluorescence signal was acquired using a Fluoromax-4 photon counting spectrofluorometer (Horiba-Jobin Yvon), and was monitored at 25°C, first with just the hybridization buffer, then with the molecular beacon, and lastly with the corresponding complementary DNA oligonucleotide target. It was recorded every 2 sec, with an integration occurring over 0.5 sec. The excitation and emission slit widths were each 2 nm, with excitation/emission wavelengths of 649/670 nm. The signal-to-background ratio was calculated by dividing the maximum fluorescence signal (CPS) by the difference between the fluorescence signal of the molecular beacon alone and the buffer background.

*In vitro* transcription of full-length *osk* RNA, and hybridization reactions with molecular beacons and *osk* RNA were performed as previously described (Catrina et al. 2012). Briefly, the hybridization reaction was performed in hybridization buffer (60 µl total volume: 80nM molecular beacon was incubated with 60nM folded *osk* RNA), and the change in fluorescence signal was recorded at 25°C every 10 sec, with 2 sec integration time and using 3 nm and 2 nm excitation (649 nm) and emission (670 nm) slit widths, respectively. The RNA was folded in hybridization buffer two ways: 1) incubation at 95 ºC for 10 sec and immediately placed on ice, and 2) incubation at 70 ºC for 10 min, followed by slow cooling over 30 min to 30 °C and placed on ice until ready to use. The sense strand of full-length *osk* mRNA was *in vitro* transcribed using a T7MegaScript Ambion Kit (ThermoFisher-Ambion), following the manufacturer’s instructions using a PCR fragment amplified from a plasmid containing *osk* cDNA inserted into a pBluescript KS+ vector (kind gift from R. Lehmann, NYU Medical School). After the RNA hybridization reaction reached a first plateau (for osk82 and osk2216) we added 3 μg of complementary DNA oligonucleotide and continued the acquisition until a maximum plateau was reached. For DNA hybridization analysis, the raw fluorescence signal (CPS) was normalized the maximum fluorescence signal, and data was fitted using non-linear regression for binding kinetics of one phase association (GraphPad Prism). For RNA hybridization analysis, the raw fluorescence signal (CPS) was normalized the maximum fluorescence signal with target *osk* RNA using the first plateau obtained from fitting the hybridization data as described above, or using the maximum fluorescence signal when adding complementary DNA oligonucleotide to the RNA-hybridization reaction after it reached the first plateau. The graphs show representative curve profiles (Fig. 6) or nonlinear fit using GraphPad Prism (Fig. 7) from three or more independent thermal denaturation, or hybridization experiments, respectively. The average maximum fluorescence signal (CPS) from *in vitro* RNA hybridization data was calculated using GraphPad Prism using the values fitted for the first plateau, as described above.

### Live cell imaging in *D. melanogaster* oocytes and image quantification

Newly hatched Oregon R-P2 (Bloomington stock #2376) female flies were fed fresh yeast paste for 2−4 days. Whole ovaries were dissected in Halocarbon Oil 700 (Sigma-Aldrich) directly on a glass coverslip and then teased apart into individual egg chambers. After mounting the glass coverslip onto the stage of the spinning disc microscope, a solution containing 300 ng/μl (∼23 μM) molecular beacon and 30 ng/ul TMR-labeled DNA probes specific to *gurken* mRNA designed for smFISH (~22 nt in length) in hybridization buffer was microinjected into the nurse cell most proximal to the oocyte. As control for the molecular beacon quantity injected, we acquired an optical stack of 41 slices (1 µm each) for the TMR channel only and with a 40x objective (oil, NA = 1.25). Image acquisition of XYZCt stacks was performed with a 63x objective (oil, NA = 1.4) for TMR and Cy5 channels, 14 optical slices (0.3 µm each), every two minutes for up to 30 min, on a Leica DMI-4000B inverted microscope (Leica Microsystems Inc.) mounted on a TMC isolation platform (Technical Manufacturing Corporation), and equipped with a Yokogawa CSU 10 spinning disk head and Hamamatsu C9100-13 ImagEM EMCCD camera (Perkin-Elmer), a 638 nm diode laser (Spectra Services Inc.), and an Eppendorf Patchman-Femtojet microinjector (Eppendorf Inc.). The images were acquired using Volocity 6.3.0 acquisition software (Perkin-Elmer) and then processed with Fiji/ImageJ software (NIH) (Schindelin et al. 2012). For Supplemental Movie S1, after maximum intensity XY projection of Z-stack, the brightness and contrast were adjusted for all 16-bit acquisition files using values optimal for osk1236, which presented the brightest *in vivo* fluorescence signal of all three *osk* molecular beacons. For Supplemental Movie S2-S4, after maximum intensity XY projection of Z-stack, the resulting time series were brightness and contrast adjusted using optimal values for each molecular beacon.

In order to compare the detection efficiency of the three molecular beacons, we quantified the Cy5 fluorescence signal accumulated at the posterior of the oocyte at the initiation of acquisition (t1 = 0 min) and after four minutes (t3 = 4 min), for three developmental stages (stage 8, 9 and late 9). We used identical acquisition parameters (laser power, and exposure time, gain and sensitivity for high-quality camera acquisition mode) for all microinjections, and found that when using parameters necessary for a detectable fluorescence signal for osk82 and osk2216 molecular beacons, time points after 4 min presented a saturated fluorescence signal for osk1236. We cropped the posterior ROI for both time points and measured the ROI area and the Raw Intensity Density (RID) for each of the 14 optical slices at each of the two time points for 16-bit Z-stacks, using an ImageJ custom macro. We calculated the Average Optical Density (AOD_Cy5_ = RID_Cy5_/Area_ROI_Cy5_)) for the optical slice that showed the maximum RID at 4 min post-injection. To account for any delay in the initiation of the acquisition protocol, we calculated the increase in AOD_Cy5_ from t1 to t3. To correct for variations in the quantity of solution injected, we similarly calculated the AOD_TMR_ for an ROI_TMR_ selected from the TMR stack acquired at 40x, for the optical slice with the maximum RID_TMR_. To correct for differences in the development stage of the injected oocytes, we measured the oocyte length from the anterior to the posterior of each injected egg chamber, and calculated a correction factor (*f*_A_→P) by dividing this distance by the minimum distance of all measured oocytes. The final relative detection efficiency of each injection was calculated using Eq. 2. The images shown in Fig. 8 are for the microinjections that presented the highest target detection efficiency for each molecular beacon, and the white rectangles highlight the ROIs used to quantify each microinjection.

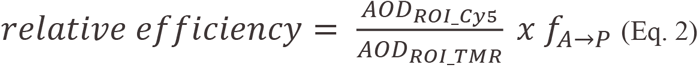

## SUPPLEMENTAL MATERIAL

Supplemental_Table_S1.pdf provides sequences, base position, GC percentage, ss-count fraction, Tm and free energies for the top 50 *osk* probes and corresponding molecular beacons designed using *PinMol* with the 31STR *mfold* input file for probe lengths ranging between 14 and 28 nt.

Supplemental_Movie_S1.avi is the time series for microinjections of osk1236, osk82 or osk2216 molecular beacon in the nurse cells of stage 8 wild type egg chambers. Images were adjusted for brightness and contrast using parameters optimal for the molecular beacon that showed the brightest signal, osk1236. Time frames acquired every 2 min for 30 min are shown as a maximum XY-projection of 14 Z-slices (0.3 µm each) at 4 frames/sec. Inset shows the maximum intensity XY-projection of the TMR injection control, 41 Z-slices, 1 μm each. Scale bar is 20 µm.

Supplemental_Movie_S2.avi is the time series for microinjections of osk1236, osk82 or osk2216 molecular beacon in the nurse cells of stage 8 wild type egg chambers. Images in each time series were adjusted for brightness and contrast using parameters optimal for the corresponding molecular beacon. Time frames acquired every 2 min for 30 min are shown as a maximum XY-projection of 14 Z-slices (0.3 µm each) at 4 frames/sec. Scale bar is 20 µm, eff = relative detection efficiency.

Supplemental_Movie_S3. avi is the time series for microinjections of osk1236, osk82 or osk2216 molecular beacon in the nurse cells of stage 9 wild type egg chambers. Images in each time series were adjusted for brightness and contrast using parameters optimal for the corresponding molecular beacon. Time frames acquired every 2 min for 30 min are shown as a maximum XY-projection of 14 Z-slices (0.3 µm each) at 4 frames/sec. Scale bar is 20 µm, eff = relative detection efficiency.

Supplemental_Movie_S4. avi is the time series for microinjections of osk1236, osk82 or osk2216 molecular beacon in the nurse cells of late stage 9, wild type egg chambers as shown in Fig. 8. Images in each time series were adjusted for brightness and contrast using parameters optimal for the corresponding molecular beacon. Time frames acquired every 2 min for 14 min (osk2216) or 30 min (osk82 and osk1236) are shown as a maximum XY-projection of 14 Z-slices (0.3 µm each) at 4 frames/sec. Scale bar is 20 µm, eff = relative detection efficiency.

## ACKNOWLEDGMENTS

We thank Ruth Lehman (Skirball Institute of Biomolecular Medicine, NYU School of Medicine) for the plasmid encoding full-length *oskar* mRNA. We thank Girish Ramrattan for useful discussions and advice on writing the *PinMol* software, and David Mathews (School of Medicine and Dentistry, University of Rochester Medical Center) for his advice and guidance with the *RNAstructure* package. We thank Salvatore A.E. Marras (Public Health Research Institute Center, Rutgers University) for the synthesis of osk82 and osk2216 molecular beacons, for the labeling and purification of the osk82, osk2216 and osk1236 molecular beacons, and for his edits, suggestions and comments on this manuscript. This work was supported by a National Science Foundation CAREER Award 1149738 and a Professional Staff Congress-CUNY Award to DPB.

